# Meta-Analysis of Public RNA Sequencing Data of Abscisic Acid-Related Abiotic Stresses in *Arabidopsis thaliana*

**DOI:** 10.1101/2023.04.17.537107

**Authors:** Mitsuo Shintani, Keita Tamura, Hidemasa Bono

## Abstract

Abiotic stresses such as drought, salinity, and cold negatively affect plant growth and crop productivity. Understanding the molecular mechanisms underlying plant responses to these stressors is essential for stress tolerance in crops. The plant hormone abscisic acid (ABA) is significantly increased upon abiotic stressors, inducing physiological responses to adapt to stress and regulate gene expression. Although many studies have examined the components of established stress signaling pathways, few have explored other unknown elements. This study aimed to identify novel stress-responsive genes in plants by performing a meta-analysis of public RNA sequencing (RNA-Seq) data on *Arabidopsis thaliana*, focusing on five ABA-related stress conditions (ABA, Salt, Dehydration, Osmotic, and Cold). The meta-analysis of 216 paired datasets from five stress conditions was conducted, and differentially expressed genes were identified by introducing a new metric, called TN (stress-treated (T) and non-treated (N))-score. We revealed that 14 genes were commonly upregulated and 8 genes were commonly downregulated across all five treatments, including some that were not previously associated with these stress responses. On the other hand, some genes regulated by salt, dehydration, and osmotic treatments were not regulated by exogenous ABA or cold stress, suggesting that they may be involved in the plant response to dehydration independent of ABA. Our meta-analysis revealed a list of candidate genes with unknown molecular mechanisms in ABA-dependent and ABA-independent stress responses. These genes could be valuable resources for selecting genome editing targets and potentially contribute to the discovery of novel stress tolerance mechanisms and pathways in plants.

## Introduction

Drought and salinity are the major abiotic stressors in plants. These stresses are becoming more severe with climate change and negatively affect crop growth, leading to yield loss (Pereira, 2016; Zhu, 2016; Mareri et al., 2022). Drought, salt stress, and secondary osmotic stress induced by them reduce plant water availability, leading to dehydration stress (Dhankher and Foyer, 2018; Ma et al., 2020). The plant hormone abscisic acid (ABA) increases significantly by these dehydration stresses and induces physiological responses to adapt to the stresses, such as stomatal closure and regulation of gene expression (Sah et al., 2016; Dar et al., 2017; Vishwakarma et al., 2017). Dehydration-responsive genes can be divided into two categories: expression regulated by ABA or specifically regulated by dehydration stress, but not by ABA (Shinozaki and Yamaguchi-Shinozaki, 2000; Yoshida et al., 2014). Additionally, ABA is synthesized under cold stress conditions (Aslam et al., 2022). The pathways that respond to cold stress and their associated gene expression share similarities with those that respond to dehydration stress (Shinozaki and Yamaguchi-Shinozaki, 2000; Yamaguchi-Shinozaki and Shinozaki, 2006). However, the expression of some genes is regulated by stimuli specific to cold stress, such as a decrease in temperature (Shinozaki and Yamaguchi-Shinozaki, 2000; Yamaguchi-Shinozaki and Shinozaki, 2006). Thus, plants respond to stress through common or uncommon pathways (Shinozaki and Yamaguchi-Shinozaki, 2000; Yamaguchi-Shinozaki and Shinozaki, 2006). Therefore, it is important to clarify the characteristics of stress-responsive genes, such as their regulatory stress and whether they are ABA-responsive (ABA-regulated or ABA-unregulated), to understand the complex regulatory mechanisms of stress responsiveness in plants. The core of the ABA signaling pathway consists of three types of proteins: ABA receptors (PYR/PYL/RCAR: pyrabactin resistance/PYR-like/regulatory component of ABA receptor), protein phosphatases (PP2C: protein phosphatases type-2C), and protein kinases (SnRK2: SNF1-related protein kinase 2) (Cutler et al., 2010; Zhang et al., 2015). These proteins regulate the activity of ABA signaling. ABA binds to receptors (PYR/PYL/RCAR) that inhibit phosphatases (PP2C) and activate kinases (SnRK2) that regulate various physiological responses through signaling cascades (Cutler et al., 2010). ABA-activated SnRK2 phosphorylates transcription factors such as ABA-RESPONSIVE EREMENT-BINDING PROTEIN (AREB)/ABRE-binding factors (ABFs) and ABA INSENSITIVE 5 (ABI5), thereby controlling the expression of downstream ABA-responsive genes (Furihata et al., 2006; Sah et al., 2016). One of the most crucial SnRK activated by ABA is Open Stomata 1 (OST1)/SRK2E/SnRK2.6, which regulates ABA-mediated stomatal closure (Mustilli et al., 2002; Yoshida et al., 2002; Lee et al., 2009). In addition, recent studies have revealed a crucial role for B2 and B3 Raf-like kinases (RAFs) as upstream regulators in the core ABA signaling pathway (Lin et al., 2020; Takahashi et al., 2020). These RAFs are required for ABA signaling for the activation of SnRK2 proteins, particularly OST1/SRK2E/SnRK2.6 (Takahashi et al., 2020; Lin et al., 2021). Furthermore, in the downstream signaling of ABA, several genes play pivotal roles. For instance, *RESPONSIVE TO DESICCATION 22 (RD22)* is recognized as a representative ABA-responsive gene (Abe et al., 1997). Moreover, genes such as the *low-temperature-responsive protein 78 (LTI78)/desiccation-responsive protein 29A (RD29A)* is induced by both ABA-dependent and ABA-independent pathways under conditions of drought, salinity, and cold (Yamaguchi-Shinozaki and Shinozaki, 1993, 1994). Also, genes located downstream of transcription factors like dehydration-responsive element binding 1s (DREB1s/CBFs) and DREB2s are known to be induced through pathways independent of ABA regulation, contributing to responses against cold, salt, and dehydration stresses(Yamaguchi-Shinozaki and Shinozaki, 1994, 2006; Liu et al., 1998; Lata and Prasad, 2011). Although several studies have examined the core elements of these signals, few have explored other novel elements. The data-driven studies have the advantage of analyzing large and independent datasets, which can lead to the identification of novel targets, distinct from the extensively studied established factors and accelerate the development of stress-tolerant crops (Bono, 2021; Ono and Bono, 2021; Tamura and Bono, 2022).

This study aimed to identify novel stress-responsive genes in plants by performing a meta-analysis of publicly available RNA sequencing (RNA-Seq) data from five ABA-related stress conditions (ABA, Salt, Dehydration, Osmotic, and Cold) in *Arabidopsis thaliana* with a focus on ABA, a factor common to many stresses. The meta-analysis revealed that 14 genes were commonly upregulated and 8 genes were commonly downregulated across all five treatments, including some that were not previously associated with these stress responses. However, some genes regulated by salt, dehydration, and osmotic treatments were not regulated by exogenous ABA or cold stress, suggesting that they may be involved in the plant response to dehydration independent of ABA.

## Materials and Methods

### Curation of public gene expression data

RNA-Seq data relevant to stress conditions involving abscisic acid were collected from the public database, National Center for Biotechnology Information Gene Expression Omnibus (NCBI GEO) (Barrett et al., 2013). A comprehensive search in GEO using the search query: (“ABA“[All Fields] OR “abscisic acid”[All Fields] OR “salt”[All Fields] OR “NaCl”[All Fields] OR “salinity”[All Fields] OR “dehydration”[All Fields] OR “osmotic”[All Fields] OR “mannitol”[All Fields] OR “drought”[All Fields] OR “cold”[All Fields] OR “low temperature”[All Fields]) AND “*Arabidopsis thaliana*”[porgn] AND “Expression profiling by high throughput sequencing”[Filter] AND (“0001/01/01”[PDAT] : “2022/08/03”[PDAT]). As a result of manual curation, the collected data were divided into five treatments: ABA, Salt, Dehydration, Mannitol, and Cold. The numbers of pairs of treated and control samples were 90, 53, 27, 12, and 34, respectively. In total, 216 paired datasets collected in this manner were used for the meta-analysis. In this study, a “paired dataset” refers to a set of two samples: one subjected to a specific stress treatment (such as abscisic acid, salt, dehydration, mannitol, or low temperature) and a corresponding control sample not subjected to the stress treatment. These paired datasets provide the basis for comparing gene expression changes induced by these stress conditions. The RNA-Seq data used in this meta-analysis are summarized in the table available online (Supplementary Table 1; https://doi.org/10.6084/m9.figshare.22566583.v6).

### Gene expression quantification

We used the SRA Toolkit (v3.0.0) (https://github.com/ncbi/sra-tools (Accessed Sep 12, 2023)) to retrieve FASTQ-formatted files for each RNA-Seq run accession number, with prefetch and fastq-dump commands, and then concatenated the files for the same experiments. Quality control of the raw reads was performed using Trim Galore (v0.6.7) (https://github.com/FelixKrueger/TrimGalore (Accessed Sep 12, 2023)) with Cutadapt (v4.1) (Martin, 2011). Transcripts were quantified using Salmon (v1.8.0) (https://github.com/COMBINE-lab/salmon (Accessed Sep 12, 2023)) (Patro et al., 2017) with a reference cDNA sequence downloaded from Ensembl Plants (release 53) (Arabidopsis_thaliana.TAIR10.cdna.all.fa.gz). As a result, the quantitative RNA-Seq data were calculated as transcripts per million (TPM). The transcript-level TPM were summarized to the gene level using tximport (v1.26.1) (https://bioconductor.org/packages/release/bioc/html/tximport.html (Accessed Sep 12, 2023)).

These operations were performed according to the workflow in SAQE (https://github.com/bonohu/SAQE (Accessed Sep 12, 2023)) (Bono, 2021). The TPM data are available online (Supplementary Table 2; https://doi.org/10.6084/m9.figshare.22566583.v6).

### Calculation score

Gene expression data from different experiments were normalized by calculating the TN-ratio, which represents the ratio of gene expression between stress-treated (T) and non-treated (N) samples.

The TN-ratio was calculated using the following equation:

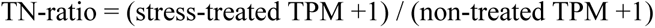

If the TN-ratio was higher than the threshold, the gene was considered upregulated; if it was less than the reciprocal of the threshold, it was deemed downregulated; otherwise, the gene was unchanged. To classify the upregulated and downregulated genes, we evaluated 2-fold, 5-fold, and 10-fold thresholds and finally chose the 2-fold threshold; therefore, genes with a TN-ratio higher than 2 were classified as upregulated, whereas genes with a TN-ratio lower than 0.5 were classified as downregulated. The TN-score of each gene was determined by subtracting the number of downregulated experiments from the number of upregulated experiments to assess the changes in gene expression under stress conditions across experiments.

TN-ratio and TN-score were calculated using a set of scripts at (https://github.com/no85j/hypoxia_code (Accessed Sep 12, 2023)) (Ono and Bono, 2021) and the data are available online (Supplementary Table 3, 4; https://doi.org/10.6084/m9.figshare.22566583.v6).

### DESeq2 analysis

To support the results with our TN-score method, we compared them with those generated using the DESeq2 package (v 1.40.2) (https://bioconductor.org/packages/release/bioc/html/DESeq2.html), a widely used tool for identifying differentially expressed genes from RNA-seq data. We set a threshold for significant expression changes at a 2-fold upregulated or downregulated (log2FoldChange ≥ |1|), applying multiple false discovery rate (FDR) thresholds (padj) of less than 0.05, 0.01, 0.005, and 0.001 to identify statistically significant changes. The analysis was conducted with ABA-treated and untreated samples using read count data. In subsequent analysis, we used the FDR thresholds of less than 0.05 and 0.001. The read count data and DESeq2 results are available online (Supplementary Table 5, 6; https://doi.org/10.6084/m9.figshare.22566583.v6).

### Gene set enrichment analysis

The web tool Metascape (https://metascape.org/ (Accessed Sep 12, 2023)) was used for gene set enrichment analysis to analyze the differentially expressed gene sets. The analysis was based on a queried gene list and the corresponding terms and p-values were examined.

### Visualization

We used a web-based Venn diagram tool (https://bioinformatics.psb.ugent.be/webtools/Venn/ (Accessed Sep 12, 2023)) and the UpSet tool (https://asntech.shinyapps.io/intervene/ (Accessed Sep 12, 2023)) to search for and visualize overlapping genes.

## Results

### Data curation of RNA-Seq from public database

RNA-Seq data on abscisic acid (ABA) and stress conditions were collected from the public database NCBI Gene Expression Omnibus (GEO). A comprehensive keyword search yielded a list of experimental data series. In contrast to microarray data from different platforms, RNA-Seq data, mostly from Illumina, are suitable for comparative analyses among studies by different research groups. Therefore, only the RNA-Seq data were used in this study. Moreover, *Arabidopsis* data were selected for the analysis because of the abundance of data in the database compared with other plants. No mutant or transgenic lines were selected and wild type samples were used. Abscisic acid, salt (NaCl), dehydration, osmotic (mannitol), and low temperature (4 °C or 10 °C) were used as stress treatment conditions. The data were curated as 216 paired datasets of stress-treated and control samples. A total of 216 pairs were used in the meta-analysis, and the tissue breakdown was as follows: 139 seedlings, 28 leaves, 23 rosette leaves, 14 roots, 6 shoots, and 6 bud tissues (Figure 1). Metadata for the curated datasets, including Sequence Read Archive (SRA) study ID, run ID, sample tissue, treatment type, treatment time, and sequence library type, are available on Supplementary Table 1.

**Figure 1.**
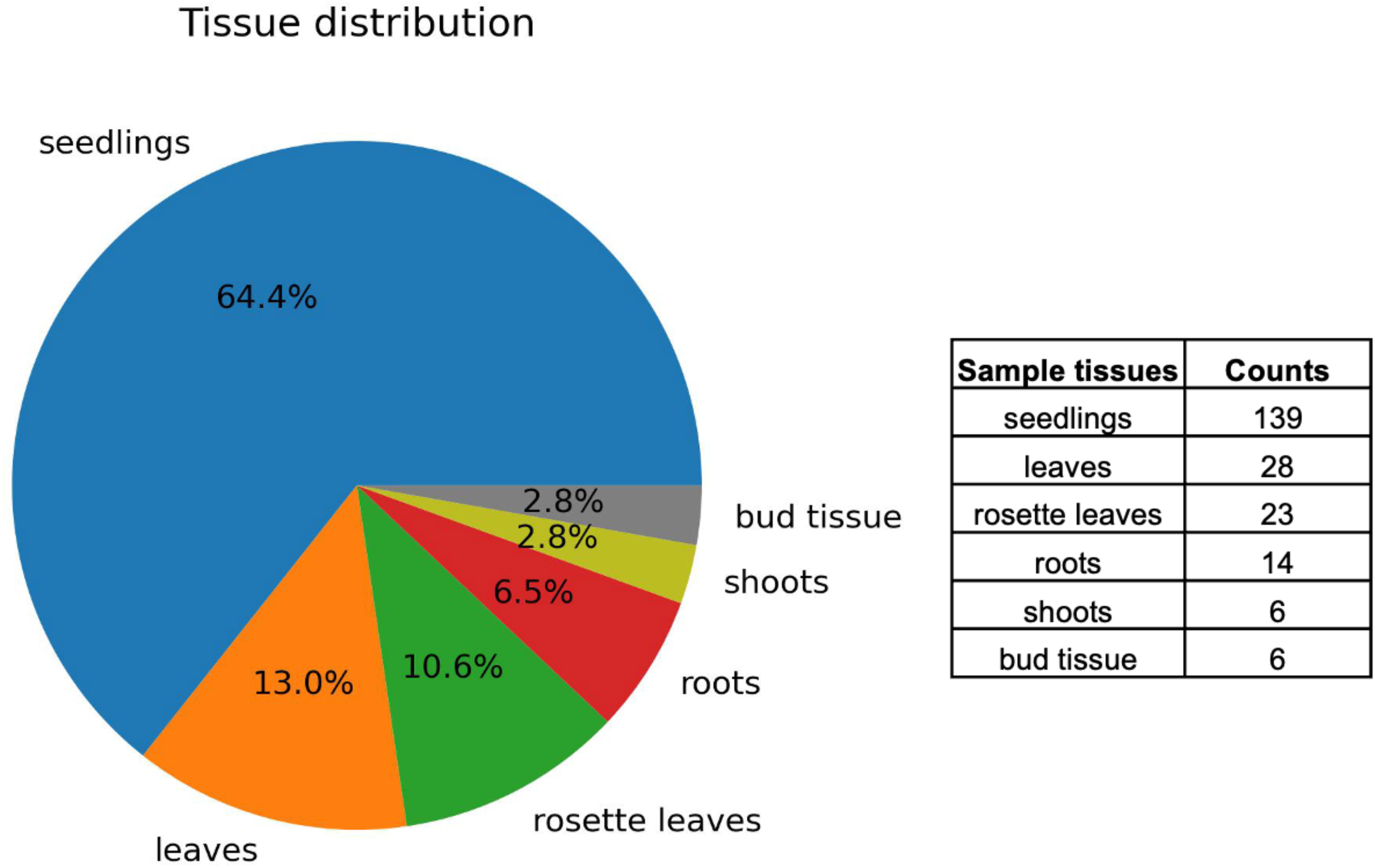
Distribution of tissue types in curated RNA-Seq data. Visualization of 216 paired datasets showing the distribution of tissues such as seedlings, leaves, rosette leaves, roots, shoots, and bud tissues.

### Identification of differentially expressed genes

Differentially expressed genes using RNA-Seq data in *Arabidopsis* under each stress condition were meta-analyzed by calculating the TN (stress-treated (T) and non-treated (N))-ratio and the TN-score of each gene. After considering various thresholds, we selected a 2-fold threshold (TN2). This threshold was slightly lower to provide a comprehensive analysis. More severe scores for the 5-fold (TN5) and 10-fold (TN10) thresholds were also calculated and are listed in the table. The lists of upregulated and downregulated genes are available online with the results of multiple thresholds considered (Supplementary Table 7, 8; https://doi.org/10.6084/m9.figshare.22566583.v6). The differences between the numbers of upregulated and downregulated experiments were calculated as the TN-scores for each gene. Thus, a higher score indicated a trend toward increased expression over the entire experiment, whereas a lower (negative) score indicated a trend toward decreased expression over the entire experiment. In the meta-analysis, we defined upregulated and downregulated as approximately top 500 genes with the highest and lowest TN2 scores, respectively. The ranges of TN2 scores for the upregulated and downregulated genes are summarized in Table 1.

**Table 1.** The ranges of TN2 scores for the upregulated and downregulated genes.

| **Treatment**<br>**Type** | **Up or Down** | **TN-Score (S)** | **Number of Genes** |
| --- | --- | --- | --- |
| ABA | upregulated | $47 \leq S \leq 88$ | 489 |
| ABA | downregulated | $-81 \leq S \leq -31$ | 486 |
| Salt | upregulated | $18 \leq S \leq 41$ | 501 |
| Salt | downregulated | $-36 \leq S \leq -11$ | 446 |
| Dehydration | upregulated | $12 \leq S \leq 24$ | 457 |
| Dehydration | downregulated | $-17 \leq S \leq -9$ | 431 |
| Mannitol | upregulated | $7 \leq S \leq 12$ | 520 |
| Mannitol | downregulated | $-11 \leq S \leq -6$ | 532 |
| Cold | upregulated | $16 \leq S \leq 32$ | 474 |
| Cold | downregulated | $-32 \leq S \leq -14$ | 551 |

Similarly, for datasets comprising pairs of ABA-treated and untreated samples, we conducted analyses using DESeq2 with read count data. In the DESeq2 analysis, we considered four FDR (adjusted p-value, padj) thresholds: 0.05, 0.01, 0.005, and 0.001. We present the overlap between the lists of genes identified as upregulated or downregulated at the highest (FDR < 0.05) and lowest (FDR < 0.001) thresholds, and the list of genes obtained using the TN2 score, in UpSet plots (Supplementary Figure 1). Among the 489 genes selected based on the TN2 score, 468 were overlapped with the genes suggested to be significantly upregulated by DESeq2 analysis at FDR < 0.05, and 464 at FDR < 0.001. For the downregulated genes, out of the 486 genes selected by the TN2 score, 320 overlapped with those suggested to be significantly downregulated by DESeq2 at FDR < 0.05, and 293 at FDR < 0.001. The results of the DESeq2 analysis and the UpSet plots are accessible online (Supplementary Table 6 and Supplementary Figure 1; https://doi.org/10.6084/m9.figshare.22566583.v6).

### Enrichment analysis of differently expression genes

We performed enrichment analysis using Metascape for five different treatment types of gene sets to characterize the differentially expressed genes (Supplementary Figure 2; https://doi.org/10.6084/m9.figshare.22566583.v6). The two most significantly enriched terms in each gene set from the enrichment analysis are summarized in table (Table 2). The most significantly enriched terms in the ABA, salt, and dehydration upregulated gene sets were all consistent with “GO:0009651 response to salt stress“. Focusing on the downregulated genes, the light-responsive genes annotated as “GO:0009642 response to light intensity” were commonly enriched in dehydration and cold treatment. The auxin-responsive genes with “GO:0009733 response to auxin” were commonly enriched in dehydration and mannitol treatment. The lists of all genes and the results of enrichment analysis are available online (Supplementary Table 7, 8; https://doi.org/10.6084/m9.figshare.22566583.v6).

**Table 2.** The two most significantly enriched terms in each gene set from the enrichment analysis.

| Treatment Type | Up or Down | Term | Description | LogP | Log(q-value) |
| --- | --- | --- | --- | --- | --- |
| ABA | upregulated | GO:0009651 | response to salt stress | -28.758572 | -24.99 |
| ABA | upregulated | GO:0071215 | cellular response to abscisic acid stimulus | -20.943027 | -17.652 |
| ABA | downregulated | GO:0050832 | defense response to fungus | -14.485652 | -10.717 |
| ABA | downregulated | GO:0001666 | response to hypoxia | -7.9936821 | -4.568 |
| Salt | upregulated | GO:0009651 | response to salt stress | -31.120518 | -27.352 |
| Salt | upregulated | GO:0009753 | response to jasmonic acid | -18.821332 | -15.402 |
| Salt | downregulated | GO:0001666 | response to hypoxia | -10.473446 | -6.886 |
| Salt | downregulated | GO:0009733 | response to auxin | -6.5022334 | -3.579 |
| Dehydration | upregulated | GO:0009651 | response to salt stress | -36.0593 | -32.291 |
| Dehydration | upregulated | GO:0010243 | response to organonitrogen compound | -19.3438 | -15.876 |
| Dehydration | downregulated | GO:0009642 | response to light intensity | -19.290212 | -15.522 |
| Dehydration | downregulated | GO:0009733 | response to auxin | -17.262779 | -13.795 |
| Mannitol | upregulated | GO:0009751 | response to salicylic acid | -28.659664 | -24.921 |
| Mannitol | upregulated | GO:0090693 | plant organ senescence | -26.790069 | -23.499 |
| Mannitol | downregulated | GO:0033013 | tetrapyrrole metabolic process | -13.988496 | -10.22 |
| Mannitol | downregulated | GO:0009733 | response to auxin | -7.0652901 | -3.774 |
| Cold | upregulated | GO:0009409 | response to cold | -24.0776 | -20.309 |
| Cold | upregulated | GO:0042254 | ribosome biogenesis | -20.3946 | -16.927 |
| Cold | downregulated | GO:0009642 | response to light intensity | -31.842859 | -28.123 |
| Cold | downregulated | ath00196 | Photosynthesis - antenna proteins - *Arabidopsis thaliana* (thale cress) | -15.580344 | -12.162 |

In addition, enrichment analysis was performed on all genes whose expression was suggested to be upregulated by DESeq2 analysis (FDR < 0.05; 1144 genes, Supplementary Figure 3A; FDR < 0.001; 1018 genes, Supplementary Figure 3B) and those genes from which genes selected based on TN2 score were excluded (FDR < 0.05; 676 genes, Supplementary Figure 3C; FDR < 0.001; 554 genes, Supplementary Figure 3D). In the enrichment analysis for all genes selected by DESeq2, the top two terms, even with either threshold, were “GO:0009651 response to salt stress” and “GO:0071215 cellular response to abscisic acid stimulus” (Supplementary Figure 3A, 3B). This is consistent with the results of the enrichment analysis for the genes selected based on the TN2 score. We also performed enrichment analysis on genes exclusively selected by DESeq2 analysis, excluding those selected based on the TN2 score (Supplementary Figure 3C, 3D). Despite the number of genes being nearly the same as those selected by the TN2 score, the results did not include “GO:0071215 cellular response to abscisic acid stimulus,” and the enrichment of significant terms was not as pronounced as observed in the enrichment analysis of genes selected based on the TN2 score. The lists of all genes and the results of enrichment analysis are available online (Supplementary Table 9 and Supplementary Figure 3; https://doi.org/10.6084/m9.figshare.22566583.v6).

### Exploring commonly regulated genes by three treatments: ABA, salt, dehydration

A previous study demonstrated that some genes are involved in the regulation and response to multiple stresses and that these genes have a potential role in universal stress tolerance in plants (Husaini, 2022). We investigated whether such candidate genes were differentially expressed under three stress conditions: ABA, salt, and dehydration. While many genes were commonly regulated under the three conditions, we identified 166 upregulated (ABA_Up Salt_Up Dehydration_Up) and 66 downregulated genes (ABA_Down Salt_Down Dehydration_Down) under the three different conditions (Figure 2). The meta-analysis also showed that the expression of *RELATED TO ABI3/VP1 1* (*RAV1)*, *RAV2*, *PYR1*, *PYL4*, *PYL5*, *PYL6*, and *PCAR3* (*PYL8*) was downregulated by ABA, salt, and dehydration stress, which is consistent with previous studies (Chan, 2012; Feng et al., 2014; Fu et al., 2014).

**Figure 2.**
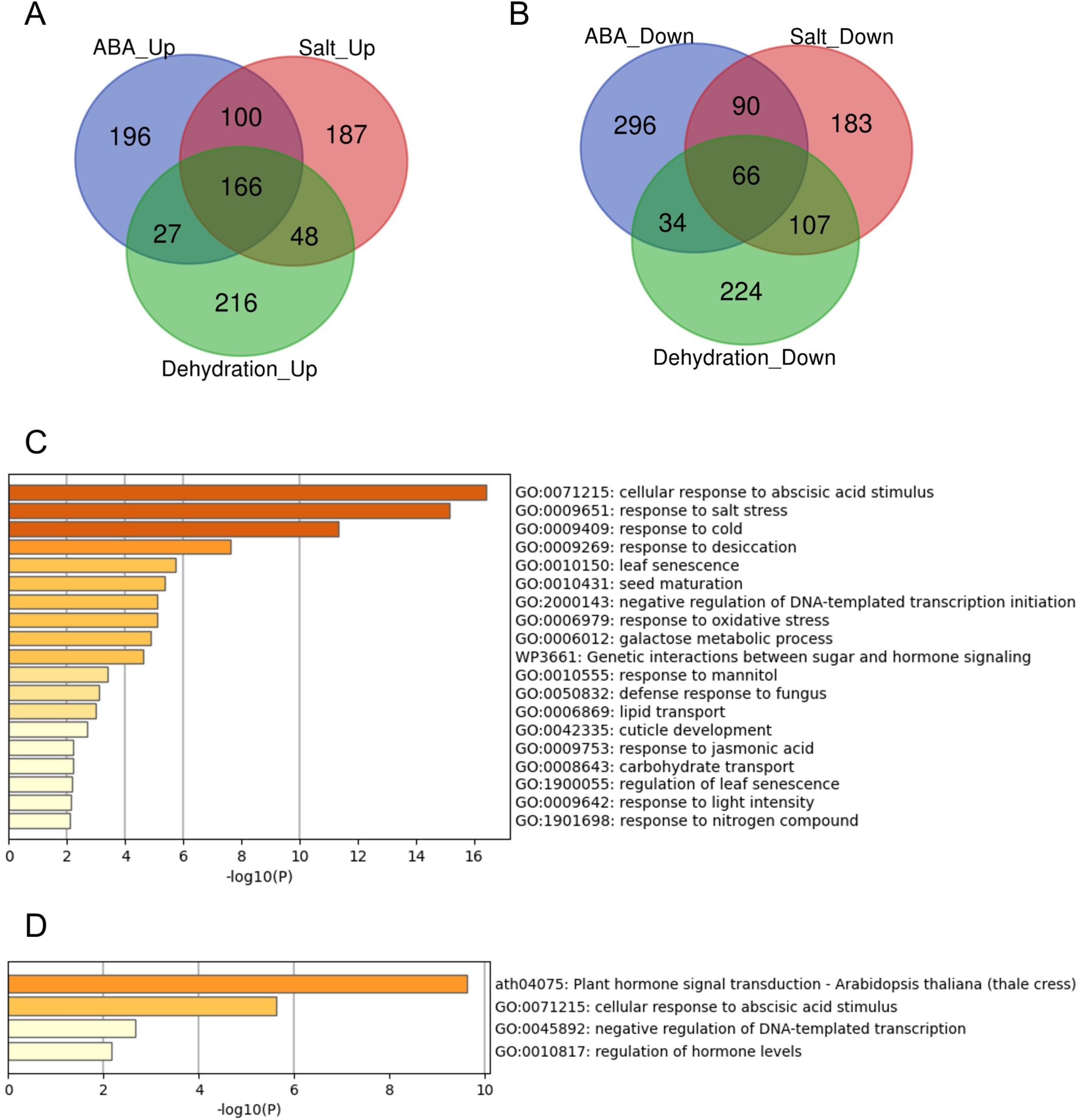
Analysis of the overlaps of upregulated or downregulated genes in 3 different types of treatments. Venn diagrams and gene set enrichment analysis in (A-D) represent the upregulated or downregulated genes in ABA, Salt, and Dehydration treatments. (A): Venn diagram of the upregulated genes, (B): Venn diagram of the downregulated genes, (C): Gene set enrichment analysis of the commonly upregulated genes, (D): Gene set enrichment analysis of the commonly downregulated genes.

Furthermore, we performed enrichment analysis of these commonly regulated genes to evaluate gene characteristics. The results showed that the top two significant enrichment terms for the 166 commonly upregulated genes (ABA_Up Salt_Up Dehydration_Up) were “GO:0071215 cellular response to abscisic acid stimulus” and “GO:0009651 response to salt stress”. In addition, the top two significant enrichment terms for the 66 commonly downregulated genes (ABA_Down Salt_Down Dehydration_Down) were “ath04075 Plant hormone signal transduction - *Arabidopsis thaliana* (thale cress)” and “GO:0071215 cellular response to abscisic acid stimulus“. The gene sets common to the three treatments were significantly enriched in terms of ABA and salt stress, indicating that these genes are likely involved in ABA and salt stress. The lists of all genes overlapping in the three treatments, and the results of the enrichment analysis are available online (Supplementary Table 10, 11; https://doi.org/10.6084/m9.figshare.22566583.v6).

### Exploring commonly regulated genes by five treatments: ABA, salt, dehydration, osmotic, cold

In the previous section, we identified commonly regulated genes across ABA, salt, and dehydration treatments. We then added mannitol and cold treatments to analyze gene overlap. The overlap of genes commonly regulated by the five treatments (ABA, salt, dehydration, osmotic stress, and cold stress) was visualized using UpSet plots and Venn diagrams (Figure 3). As a result, 14 commonly upregulated genes (ABA_Up Salt_Up Dehydration_Up Mannitol_Up Cold_Up) and 8 commonly downregulated genes (ABA_Down Salt_Down Dehydration_Down Mannitol_Down Cold_Down) were identified. The genes included in these two groups are listed in Tables 3 and 4 respectively. The genes commonly upregulated in the five treatments contained *LTI78/RD29A*, *LTI30*, *Stress-responsive protein/Stress-induced protein 1* (*KIN1)*, *COLD-REGULATED 47* (*COR47)*, and *Alcohol Dehydrogenase 1* (*ADH1)*, which are reported to be induced by ABA, salt, dehydration, and cold stress conditions (Kurkela and Franck, 1990; Jarillo et al., 1993; Yamaguchi-Shinozaki and Shinozaki, 1994; Ishitani et al., 1998; Shi et al., 2015). On the other hand, genes such as *Arabidopsis thaliana heavy metal associated domain containing gene 1* (*ATHMAD1)*, which have not been reported to have ABA-related or stress response of functions, were also identified. These genes have been identified as novel stress-responsive genes involved in the ABA response. Also, the ABA-independent pathway, which is specifically induced by osmotic stress has been reported in plants (Nakashima et al., 2014; Yoshida et al., 2014). This pathway is hypothesized to be activated independently of ABA and its closely related cold stimulus (Yamaguchi-Shinozaki and Shinozaki, 2006). Therefore, we examined three treatments, excluding ABA and cold, to explore genes involved in the ABA-independent osmotic signaling pathway. As a result, 7 commonly upregulated genes (Salt_Up Dehydration_Up Mannitol_Up) and 48 commonly downregulated genes (Salt_Down Dehydration_Down Mannitol_Down), regulated by salt, dehydration, and mannitol, were also identified. Notably, these genes were not included among those suggested to be regulated by ABA and cold stress in the present study, and the effects of ABA and cold on the regulation of their expression may not be significant. Therefore, they may be involved in the ABA-independent stress response pathways. The genes included in these two groups are listed in Tables 5 and 6. The lists of all genes that overlap in the five treatments are available online (Supplementary Table 12, 13; https://doi.org/10.6084/m9.figshare.22566583.v6).

**Figure 3.**
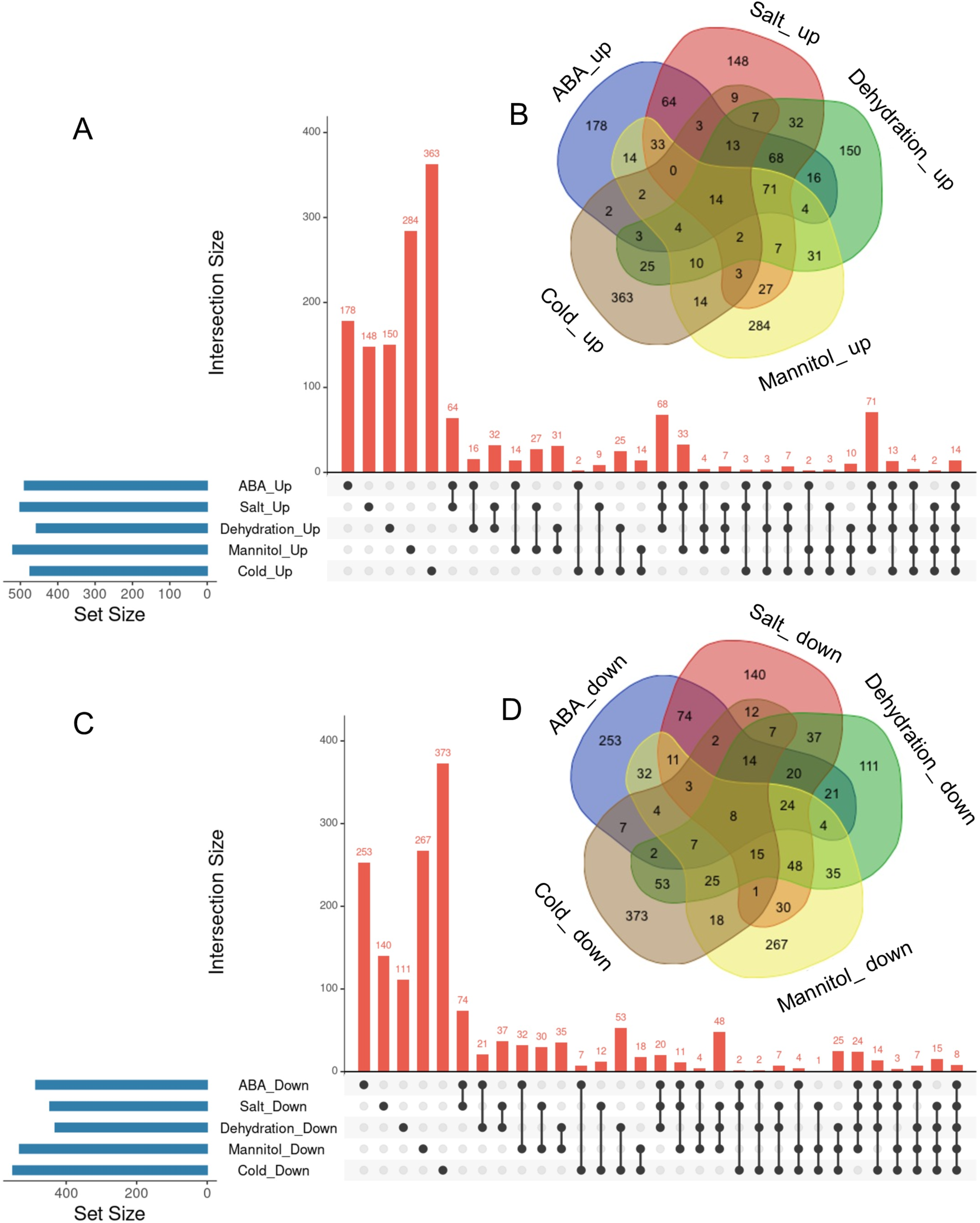
Analysis of the overlaps of upregulated or downregulated genes in 5 different types of treatments. UpSet plots and venn diagrams in (A-D) represent the upregulated or downregulated genes in ABA, Salt, Dehydration, Mannitol, and Cold treatments. (A): UpSet plots of the upregulated genes, (B): Venn diagram of the upregulated genes, (C): UpSet plots of the downregulated genes, (D): Venn diagram of the downregulated genes.

**Table 3.** The lists of commonly upregulated genes in ABA, Salt, Dehydration, Mannitol, and Cold treatments.

| Gene ID | Gene Name | Description | Regulated by Hypoxia<br>in previous meta-analysis |
| --- | --- | --- | --- |
| AT1G11210 | - | cotton fiber protein, putative (DUF761) | No |
| AT1G16850 | - | transmembrane protein | No |
| AT1G73810 | - | Core-2/I-branching beta-1,6-N-acetylglucosaminyltransferase family protein | No |
| AT4G01985 | - | hypothetical protein | No |
| AT5G11110 | SPS2F | sucrose phosphate synthase 2F | No |
| AT1G51090 | ATHMAD1 | Heavy metal transport/detoxification<br>superfamily protein | No |
| AT5G17460 | DFR1 | glutamyl-tRNA (Gln) amidotransferase subunit<br>C | No |
| AT1G67360 | LDAP1 | Rubber elongation factor protein (REF) | Induced |
| AT1G77120 | ADH1 | alcohol dehydrogenase 1 | Induced |
| AT1G20440 | COR47 | cold-regulated 47 | No |
| AT5G15960 | KIN1 | stress-responsive protein (KIN1) / stress-induced protein (KIN1) | No |
| AT1G01470 | LEA14 | Late embryogenesis abundant protein | No |
| AT3G50970 | LTI30 | dehydrin family protein | No |
| AT5G52310 | RD29A | low-temperature-responsive protein 78 (LTI78)<br>/ desiccation-responsive protein 29A (RD29A) | No |

**Table 4.** The lists of commonly downregulated genes in ABA, Salt, Dehydration, Mannitol, and Cold treatments.

| Gene ID | Gene Name | Description | Regulated by Hypoxia in previous meta-analysis |
| --- | --- | --- | --- |
| AT3G23880 | - | F-box and associated interaction domains-containing protein | No |
| AT5G19190 | - | hypothetical protein | No |
| AT4G38850 | SAUR15 | SAUR-like auxin-responsive protein family | No |
| AT4G38860 | SAUR16 | SAUR-like auxin-responsive protein family | Repressed |
| AT1G29430 | SAUR62 | SAUR-like auxin-responsive protein family | Repressed |
| AT1G29440 | SAUR63 | SAUR-like auxin-responsive protein family | Repressed |
| AT1G74670 | GASA6 | Gibberellin-regulated family protein | Repressed |
| AT4G36570 | RAD-LIKE 3 | RAD-like 3 | Repressed |

**Table 5.** The lists of commonly upregulated genes in Salt, Dehydration, and Mannitol treatments.

| Gene ID | Gene Name | Description | Regulated by Hypoxia in previous meta-analysis |
| --- | --- | --- | --- |
| AT1G17830 | - | hypothetical protein (DUF789) | Induced |
| AT1G76590 | - | PLATZ transcription factor family protein | No |
| AT5G42050 | - | DCD (Development and Cell Death) domain protein | No |
| AT3G22370 | AOX1A | alternative oxidase 1A | No |
| AT2G21590 | APL4 | Glucose-1-phosphate adenylyltransferase family protein | No |
| AT1G02850 | BGLU11 | beta glucosidase 11 | No |
| AT4G37370 | CYP81D8 | cytochrome P450, family 81, subfamily D, polypeptide 8 | No |

**Table 6.**
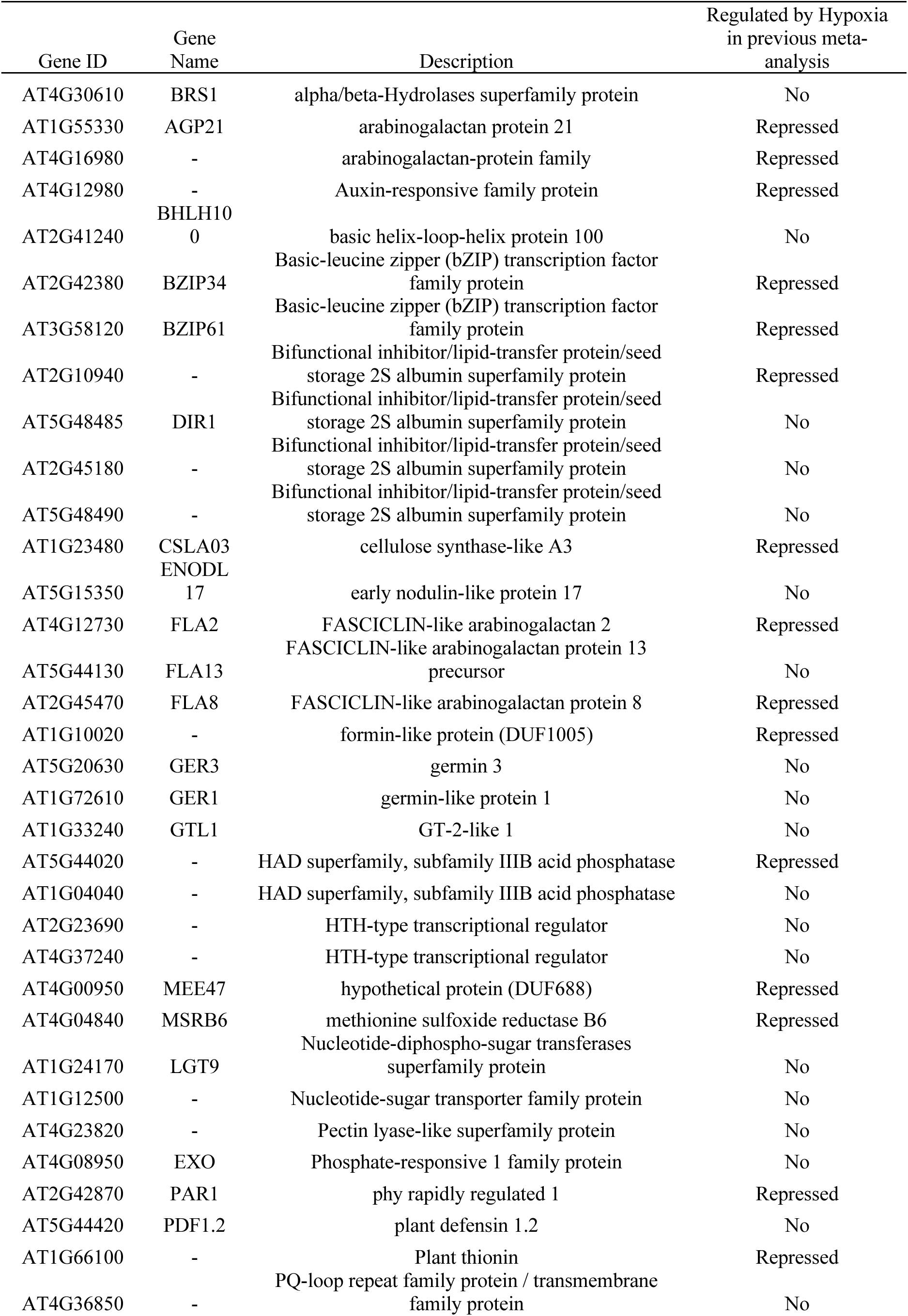
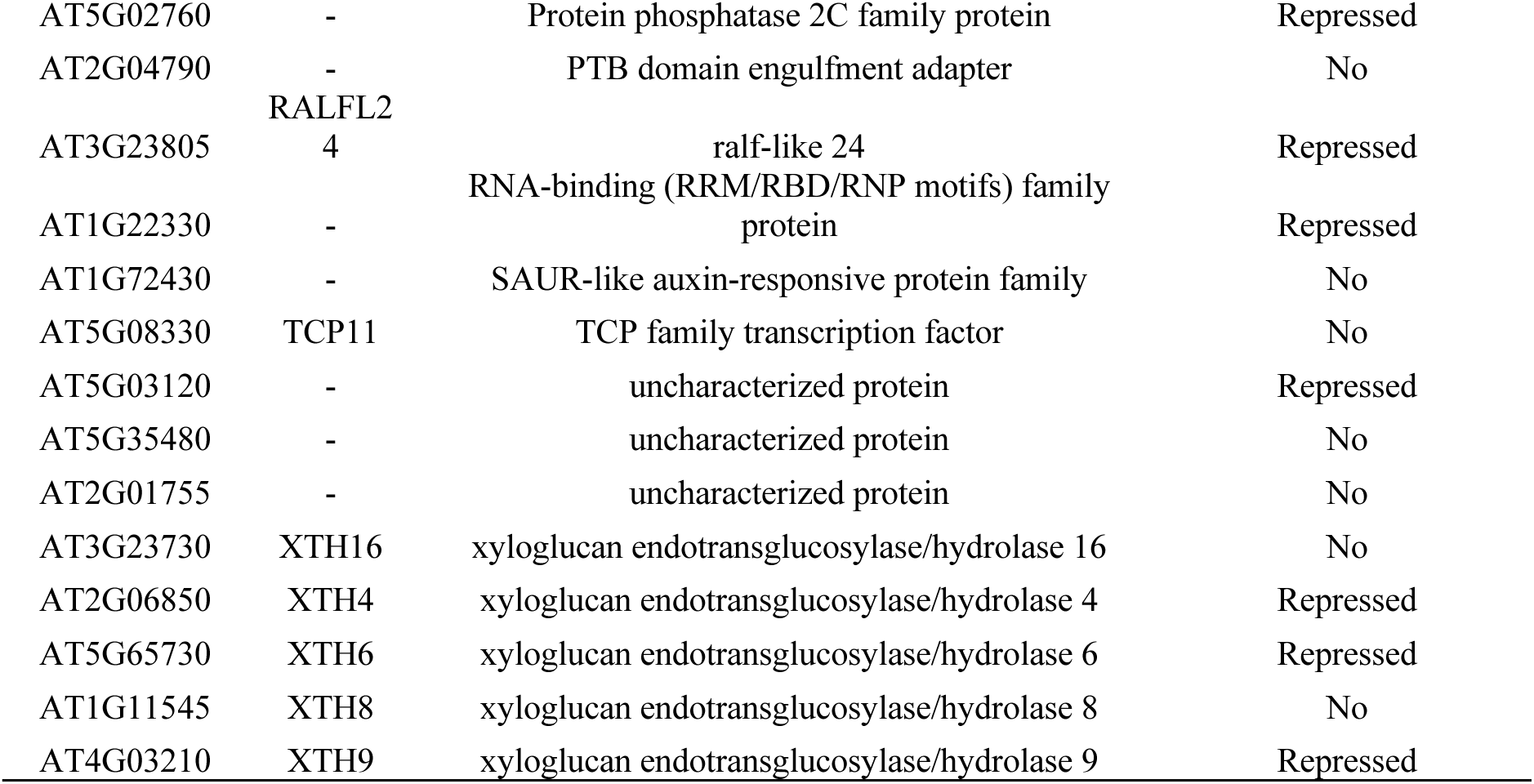
The lists of commonly downregulated genes in Salt, Dehydration, and Mannitol treatments.

## Discussion

In this study, we aimed to explore uncharacterized differentially expressed genes by meta-analyzing public RNA-Seq data of *Arabidopsis thaliana* to gain new insights into the effects of abscisic acid (ABA) and various stress conditions on gene expression. The data were collected from the public database NCBI GEO, and the study focused on RNA-Seq data because of their suitability for comparative analyses among different studies. We analyzed the effects of ABA, salt, dehydration, osmotic, and low-temperature stress on gene expression. In total, 216 paired datasets of stress-treated and control samples were used for the meta-analysis. The central to this research is the notion that by employing a comprehensive integrated analysis using large-scale data from multiple research groups, we can uncover patterns and insights that might be overlooked in individual studies. We employed the TN-score to identify differentially expressed genes under each stress condition. We defined upregulated and downregulated genes as the top 500 genes with the highest and lowest TN-scores, respectively, in the meta-analysis. Enrichment analysis was performed using Metascape to characterize the differentially expressed genes. The most significantly enriched terms in the downregulated genes, the light-responsive genes annotated as “GO:0009642 response to light intensity” were commonly enriched in dehydration and cold treatment, and the auxin-responsive genes annotated as “GO:0009733 response to auxin” were commonly enriched in dehydration and mannitol treatment. Abiotic stress negatively affects various aspects of plant photosynthesis including stomatal conductance, oxidative stress, RUBISCO activity, photosystems (PS I and PS II), photosynthetic electron transport, and chlorophyll biosynthesis (Sharma et al., 2020). These factors reduce the photosynthetic efficiency and plant growth. Therefore, it is possible that dehydration and cold treatments could have downregulated the expression of light-responsive genes “GO:0009642 response to light intensity” due to adjustments in light sensitivity and photosynthetic responses during the process of adapting to stress. In addition, osmotic stress reduces the biosynthesis of auxin, a plant growth hormone, by directly or indirectly inhibiting the auxin pathway (Naser and Shani, 2016). This might contribute to reduced plant growth under stress conditions. Auxin biosynthesis may be suppressed by osmotic stress conditions, resulting in downregulated auxin-responsive genes “GO:0009733 response to auxin“. To support the selection results of the TN2 score, we performed a comparison of genes suggested to be upregulated and downregulated by DESeq2 analysis in ABA-treated samples and those selected based on the TN2 score. The number of genes detected as unregulated by TN2 score alone was 489, and those detected as upregulated by both DESeq2 (FDR < 0.05) and TN2 scores was 468, with an overlap ratio of 95.7% (Supplementary Figure 1A).

Furthermore, a comparison of enrichment analysis results showed that the top two GO terms for all genes detected by DESeq2 and genes selected by the TN2 score were similar (Supplementary Figure 3A, 3B). However, the enrichment analysis of genes detected only by DESeq2 did not show a significant enrichment of key terms compared to those selected by the TN2 score (Supplementary Figure 3C, 3D). These suggest that employing TN-scores for gene selection not only the genes detected by commonly used tools like DESeq2 but also effectively refines the overall pool of selected genes. This indicates that utilizing TN-scores is an effective method for meta-analysis. Next, we identified genes that were commonly regulated by ABA, salt, and dehydration stress. We also performed enrichment analysis of genes commonly regulated by different stresses and verified their consistency with previously reported gene expression patterns. We expanded this analysis to include genes commonly regulated by all five treatments (ABA, salt, dehydration, osmotic, and cold). Several genes were upregulated or downregulated in response to these stress conditions, suggesting their potential roles in global plant stress tolerance mechanisms.

Fourteen upregulated genes were found to be closely involved in ABA regulation (ABA_Up Salt_Up Dehydration_Up Mannitol_Up Cold_Up, Table 3), including genes that have not been previously studied, one of which is ATHMAD1 (AT1G51090). ATHMAD1 contains a heavy metal-associated (HMA) domain that is upregulated in response to nitric oxide (NO) (Imran et al., 2016). NO plays a variety of roles in plant adaptation to abiotic stresses such as drought, salt, and cold (Fancy et al., 2017; Lau et al., 2021). NO synthesis is promoted by abscisic acid (ABA) treatment and abiotic stress, which contributes to stomatal closure during drought stress (Neill et al., 2008). Analysis of *athmad1* mutant indicated that ATHMAD1 may play a role in regulating plant growth and immunity (Imran et al., 2016). However, the function of ATHMAD1 under abiotic stress has not yet been thoroughly investigated. In this study, *ATHMAD1* expression was upregulated upon treatment with ABA, salt, dehydration, osmotic, and cold stress. Based on these findings, *ATHMAD1* expression may increase in response to NO, which is promoted by ABA treatment and abiotic stress. Therefore, analyzing the function of ATHMAD1 in plants under abiotic stress conditions is an important challenge for future research.

In addition, focusing on genes downregulated by multiple stressors, we examined eight genes considered to be deeply involved in ABA-dependent expression regulation (ABA_Down Salt_Down Dehydration_Down Mannitol_Down Cold_Down, Table 4) and 48 genes considered to be less influenced by ABA-mediated expression regulation (Salt_Down Dehydration_Down Mannitol_Down, Table 6). It was found that among the eight genes, four were *SAUR (Small Auxin Up RNA)* genes (*SAUR15, 16, 62, 63*), and among the 48 genes, five *XTH (Xyloglucan endotransglucosylase/hydrolase)* genes (*XTH4, 6, 8, 9, 16*) were included, respectively. Although SAUR and XTH gene families have distinct mechanisms and molecular targets, they play a common role in controlling plant growth and cell wall expansion. Furthermore, several studies have identified these genes during stress stimulation and changes in stress response phenotypes in mutants lacking their functions.

The *SAUR* genes are a plant-specific gene family that plays a crucial role in plant growth and development by regulating cell wall acidification in response to auxin stimulation (Ren and Gray, 2015; Stortenbeker and Bemer, 2019). Several *SAUR* genes were downregulated in response to ABA, drought, and osmotic stress. ARABIDOPSIS ZINC-FINGER 1 (AZF1) and AZF2 were induced by osmotic stress and abscisic acid, function as transcriptional repressors of 15 SAUR genes, thereby inhibiting plant growth under abiotic stress conditions (Kodaira et al., 2011). Among the genes repressed by AZF, AZF directly binds to the promoter region of *SAUR63* (Kodaira et al., 2011). SAUR63 localizes to the plasma membrane and by inhibiting plasma membrane (PM)-associated PP2C.D phosphatases, it stimulates PM H^+^-ATPase proton pump activity, thereby promoting cell growth (Nagpal et al., 2022). However, the functions of most SAUR genes under abiotic stress and their direct relationship with stress remain unclear.

XTH is an enzyme family involved in plant cell wall remodeling that catalyzes the cleavage and polymerization of xyloglucan to regulate cell elasticity and extensibility (Rose et al., 2002). XTH not only plays a role in regulating cell wall structure and morphology, but also plays a crucial role in plant adaptation to external stress (Ishida and Yokoyama, 2022). The expression of *AtXTH19* and *AtXTH23* was induced by brassinosteroids, and the *atxth19* mutant showed lower freezing tolerance than the wild type, whereas the *atxth19/atxth23* double mutant showed increased sensitivity to salt stress (Xu et al., 2020; Takahashi et al., 2021). Similarly, *AtXTH30* contributes to salt tolerance (Yan et al., 2019). Additionally, the loss-of-function of several *XTH* genes has been reported to affect metal ion tolerance. *AtXTH31* is associated with cell wall aluminum binding capacity, and mutation of this gene leads to increased aluminum tolerance (Zhu et al., 2012). Similarly, *Atxth18* mutant showed reduced sensitivity to lead stress (Zheng et al., 2021). *AtXTH4* and *AtXTH9* contribute to the regulation of xylem cell expansion and secondary growth, including the production of secondary xylem and the accumulation of secondary walls. However, the function of stress conditions has not yet been clarified (Kushwah et al., 2020). These studies suggested that XTH plays an important role in various stress responses. Therefore, understanding the functions of the five XTH genes (*XTH4, 6, 8, 9, 16*), the genes that were downregulated by ABA-independent stress pathways in this study, is a future challenge. As discussed above, the literature information on these mutant genes reinforced the correctness of our analysis.

Finally, we investigated whether the hypoxia-responsive genes identified in a previous meta-analysis in *Arabidopsis thaliana* were included in the genes we focused on in this study (Tamura and Bono, 2022). Under hypoxic conditions, root water transport is inhibited and stomatal closure is induced. This process requires aquaporins, which are proteins involved in water transport, as well as the hormones ABA and ethylene (Tan et al., 2018). Furthermore, during recovery from submergence, reactive oxygen species (ROS), ABA, ethylene, respiratory burst oxidase homolog D (RBOHD), senescence-associated gene113 (SAG113), and ORESARA1 (ORE1) regulate ROS homeostasis, stomatal opening/closing, and chlorophyll degradation (Yeung et al., 2018). These studies suggest that hypoxia and ABA levels are related to each other. The analysis demonstrated that the expression of several stress-regulated genes, which were the focus of this study, was regulated by hypoxia (Tables 3–6). These results imply that there might be an undiscovered molecular cross talk between “hypoxia” and “ABA or dehydration,” which has not been frequently discussed in previous research. The overlap of genes commonly regulated by the six treatments (ABA, salt, dehydration, osmotic, cold, and hypoxia) was visualized using UpSet plots (Figure 4). These genes are regulated by a wide range of stresses and identifying their roles could contribute to further advances in our understanding of the broad mechanisms of stress tolerance in plants. The lists of all genes overlapping in the six treatments are available online (Supplementary Table 14, 15; https://doi.org/10.6084/m9.figshare.22566583.v6).

**Figure 4.**
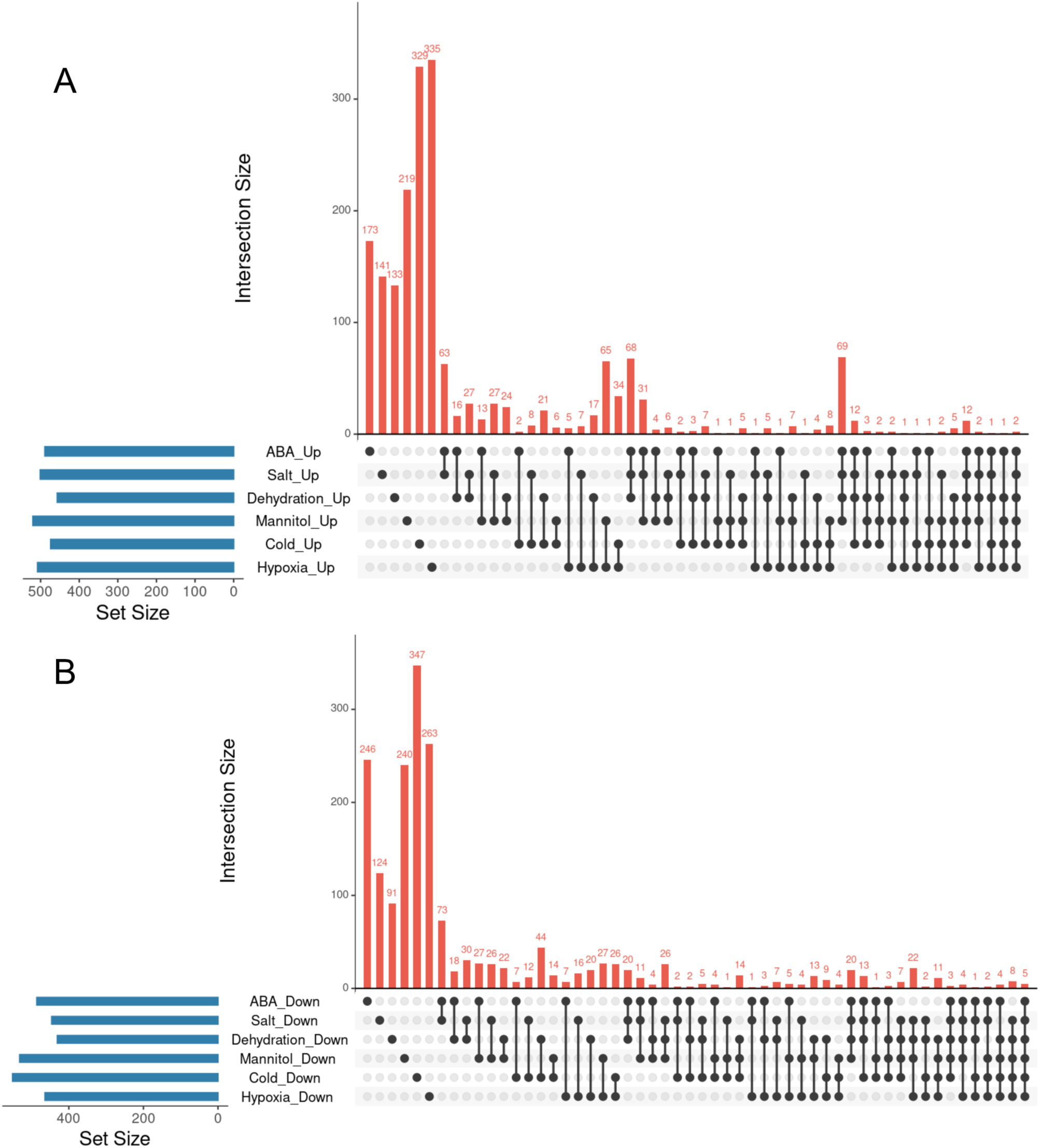
Analysis of the overlaps of upregulated or downregulated genes in 6 different types of treatments. UpSet plots in (A, B) represent the upregulated or downregulated genes in ABA, Salt, Dehydration, Mannitol, Cold, and Hypoxia treatments. (A): UpSet plots of the upregulated genes, (B): UpSet plots of the downregulated genes.

This study has several limitations that need to be considered. First, the identification of differentially expressed genes was not subjected to statistical testing, demanding caution in the interpretation of results. Second, the RNA-Seq data used in this study were collected under various growth and experimental conditions, careful consideration of these variabilities. Third, experimental validation using plant samples exposed to specific stress environments has not been performed, necessitating further research for functional clarification of these genes.

On the other hand, the strength of this study is the integration of data from multiple research projects and various stress conditions. Representative ABA-responsive or stress-responsive genes were extracted using the TN-score method. Furthermore, the candidate genes with unknown function listed in this study have the potential to provide new insights into the stress response mechanisms in plants. These findings are expected to contribute to the broadening of the future plant stress research.

In conclusion, this meta-analysis lists candidate genes that are responsive to ABA-dependent or ABA-independent stress conditions, potentially playing roles in plant responses to these stresses. In this study, we focused on the gene expression commonly found under different stresses and validated our analysis by confirming its consistency with previously reported patterns of gene expression. As the data in the database increases, it is expected that new findings will be revealed by updating and reanalyzing the datasets. Although further studies are required to reveal the role of the genes identified in this study, the list obtained in this study can be a useful decision criterion for the selection of genome editing targets and the discovery of novel molecular mechanisms. We think that our research forms the foundation for revealing patterns and insights that might have been overlooked in individual studies.

## Author Contributions

MS and HB conceived and designed the study. MS performed the experiments and collected data. KT helped the analysis. Writing—original draft: MS. All authors discussed the data and helped with manuscript preparation. HB supervised the project. All authors read and approved the final manuscript.

## Acknowledgements

Computations were performed on the computers at Hiroshima University Genome Editing Innovation Center.

## Supplementary Material

Additional data presented in this study are publicly available online (https://doi.org/10.6084/m9.figshare.22566583.v6).

